# Menstrual cycle irregularity is a biological determinant of mental health independent of sleep in adolescents

**DOI:** 10.64898/2026.08.03.742466

**Authors:** Fernanda Mayara Crispim Diogo, Lucas G. S. França, Mario André Leocadio-Miguel, Maricele Nascimento Barbosa, Carolina Virginia Macêdo de Azevedo

**Author notes:** The corresponding author: Carolina V.M. Azevedo. – Departamento de Fisiologia e Comportamento/ Universidade Federal do Rio Grande do Norte - 59078-970 Natal/RN – Brazil.

## Abstract

**INTRODUCTION:** Sex differences in mental health emerge during adolescence, a period marked by the onset and the establishment of menstrual cycle. However, is rarely examined how menstrual cycle regularity, a marker of hormonal function, modulates mental health.

**OBJECTIVE:** to analyse sex differences in mental health symptoms among adolescents considering the menstrual cycle regularity and sleep.

**METHODS:** A three-group design (female students with regular cycles/FR, n=77; with irregular cycles/FI, n=59; and male students/M, n=76) in a sample of Brazilian high-school adolescents (n=212; 14–18 years) enrolled in morning and full-time classes was used to test the hypothesis that mental health symptoms follow a graded pattern across these groups.

**RESULTS:** Mean DASS-21 scores across all groups fell at or above the Mild severity threshold for mental health subscales. GLMs confirmed a monotonic gradient increase in group order (M→FR→FI) which was associated with higher scores on all outcomes (stress β/step=4.33, p<.001; anxiety β/step=4.08, p<.001; and depression β/step=2.34, p=.010; model R²=.16, .13, .08 respectively). However, no differences were observed in sleep duration, social jetlag, chronotype, sleep quality, or sleep-debt. Then, a secondary analysis assessed sex-specific associations between socioeconomic status (SES) and mental health; higher SES was inversely related to stress, anxiety, and depression, being protective only in males (stress Males β=−2.68, p=.011/Females β=0.46, p=.614).

**CONCLUSION:** These findings support the reframing of menstrual irregularity not only as a reproductive health concern but also as a biological determinant of mental health risk in female adolescents, a vulnerability that sleep disruption and socioeconomic resources do not adequately explain.

## Introduction

Adolescence marks a critical inflection point for mental health vulnerability, with a sharp rise in stress, anxiety, and depression, which disproportionately affects females [1,2]. Epidemiological data demonstrate that the increase in mood and anxiety symptoms in female adolescents begins around puberty and intensifies through late adolescence [3]. This pattern suggests a biological, rather than purely behavioural and social dimensions, as its driving force. Despite this, most research on adolescent mental health has treated sex as a single binary covariate, rather than examining possible biological mechanisms that underlie female vulnerability, such as the menstrual cycle irregularity.

Ovarian hormones have widespread effects throughout the body, primarily influencing the central nervous system. These hormones are known to bind to receptors and to influence the activity of limbic, prefrontal and hypothalamic structures involved in stress regulation, mood and cognition [4]. Menstrual cycle irregularity, defined as cycle length consistently outside the 21–35-day range, reflects disruption of this hormonal axis and is particularly prevalent during adolescence [5,6]. Precisely, up to 50% of female adolescents report irregular cycles in the first two years following menarche [6,7]. Irregular cycles have been associated with more intense menstrual symptoms, mood imbalances, poorer sleep quality, and higher rates of psychological distress [8–10].

Nonetheless, the relationship between menstrual irregularity and mental health in adolescents remains poorly explored. Most prior work has examined irregularity as a predictor of sleep parameters, with mental health outcomes treated as downstream variables [9–11]. No published study has directly compared adolescent females with regular and irregular cycles against a male reference group. This study addresses this gap in a three-group cross-sectional design. Using male adolescents as a biological comparator and regular-cycle females as an intermediate condition, we tested the hypothesis that mental health symptoms follow a graded, monotonic pattern across the three groups: males < regular females < irregular females. We further tested whether this gradient is explained by sleep and circadian variables, and whether socioeconomic status moderates the gradient differently by sex. The study was conducted in a Brazilian public-school context with early school start times (7:00–7:30 am), a setting known to produce chronic sleep restriction in adolescents [12,13] and therefore well-suited to examining how biological vulnerability interacts with environmental stress.

## Materials and Methods

### Participants

The study was conducted with 212 high-school students (females: n = 136, age = 15.9 ± 0.9 years; males: n = 76, age = 16.1 ± 1.1 years) enrolled in either morning (classes starting at 7:00–7:30 am) or full-day classes at three public schools in the metropolitan area of Natal, Brazil (Latitude: −5.79°S, Longitude: −35.21°W). Students were recruited between 2023 and 2024, excluding school vacation periods, exam weeks, and public holidays. The female group was subdivided into female students with regular menstrual cycles (FR, n = 77) and female students with irregular menstrual cycles (FI, n = 59), based on self-report. Exclusion criteria comprised any reported sleep, psychiatric, or neurological disorder, and any current hormonal treatment, including the use of contraceptive pills. The study was approved by the Ethics Committee of the Federal University of Rio Grande do Norte (protocols 5.456.242 and 5.905.464), and informed consent was provided by the students’ legal guardians.

### Design and analytical groups

The primary analytical structure was a three-group comparison: (1) female students with regular menstrual cycles (FR), (2) female students with irregular menstrual cycles (FI), and (3) males (M). Male students served as a reference group, anchoring the lower of an expected mental health burden gradient. Regular-cycle female students served as an intermediate reference, representing the expected female hormonal baseline in the absence of cycle disruption. Irregular-cycle female students were hypothesised to show the highest mental health burden, reflecting an additional hormonal disruption beyond the baseline sex effect. Groups were compared on all sleep-wake cycle and circadian parameters and mental health symptoms. A secondary analysis examined the interaction between sex and socioeconomic status on mental health outcomes in the full sample.

### Instruments

#### Sleep and chronotype assessment

We assessed sleep and chronotype with standardised instruments. The Munich Chronotype Questionnaire (MCTQ) [14] provided weekday and weekend sleep onset and offset times, sleep duration, and chronotype (mid-sleep phase on free days corrected for sleep compensation, MSFsc). Social jetlag was computed as the discrepancy between mid-sleep phase on free days and school days [15]. Sleep quality was subjectively measured with the Pittsburgh Sleep Quality Index (PSQI) [16,17]. Scores above 5 indicate poor sleep quality. Daytime sleepiness was assessed using the Pediatric Daytime Sleepiness Scale (PDSS) [18,19]. Scores at or above 15 indicate excessive daytime sleepiness [20].

#### Mental health assessment

We applied the Depression, Anxiety and Stress Scale 21 (DASS-21) [21,22] to quantify the severity of symptoms across three distinct dimensions: stress, anxiety, and depression. Each subscale yields scores between 0 and 42, with validated severity cutoff-points (Normal, Mild, Moderate, Severe, Extremely Severe). For continuous analyses, raw subscale scores were used.

#### Menstrual cycle regularity

Cycle regularity was assessed via the Sleep Habits Questionnaire (SHQ), which includes a four-point self-report item (“very regular”, “regular”, “irregular”, “very irregular”). For the present study, responses were dichotomised: “very regular” and “regular” were classified as FR, and “irregular” and “very irregular” as FI.

#### Socioeconomic status (SES)

Finally, we assessed household socioeconomic status using the Brazilian Economic Classification Criterion [23], which assigns a continuous score based on household asset ownership and parental education. Higher scores indicate higher socioeconomic status.

### Statistical analysis

Sociodemographic variables and all sleep-wake cycle parameters were compared using one-way ANOVA for continuous and chi-square tests for categorical variables. For the primary analysis, group differences in stress, anxiety, and depression were modelled using Gaussian-identity generalised linear models (GLMs) with regularity group (reference = Male) as the focal predictor and age, standardised socioeconomic score, and school shift used as covariates. Adjusted pairwise contrasts (FR vs Male, FI vs Male, FI vs FR) were obtained using the emmeans R package [24], with Benjamini-Hochberg false-discovery-rate (FDR) correction [25] applied across the three outcomes. Effect sizes were expressed as Cohen’s *d*. The monotonic gradient hypothesis was also tested using a continuous ordered predictor (Male = 1, FR = 2, FI = 3) in the same adjusted GLM, providing an estimated β per group step. We used chi-square tests for categorical symptom severity distributions. In our secondary analysis, we examined whether the SES vs. mental health relationship differed by sex through a Gaussian-identity GLM including menstrual cycle regularity, SES, the SES × female interaction, age, school shift, and chronotype as covariates. Finally, we tested whether the mental health gradient was dependent on sleep and circadian disruption by adding group × sleep variable interaction terms to the baseline GLM. All analyses were conducted in R (version 4.5.2) [26] using emmeans [24], ggplot2 [27], and patchwork [28].

## Results

### Group comparability

Groups did not differ significantly in age, socioeconomic score, or school shift distribution. Wake-up time on weekdays differed significantly across groups (F_(2,209)_ = 4.06, p = 0.019), with male students waking later than both female groups (post-hoc comparisons: both *p* < 0.05; FR vs FI: *p* > 0.05). Daytime sleepiness also differed across groups (F_(2,209)_ = 4.83, *p* = 0.009), with both female groups reporting higher levels than male students (post-hoc comparisons: both *p* < 0.05, no difference between FR and FI groups). No significant group differences were found in sleep duration on weekdays or weekend, sleep debt, social jetlag, chronotype, or PSQI score (all all *p* > 0.23). Table 1 presents the sample characteristics for the three groups.

**Table 1.** Sample characteristics by group. Values are Mean ± SD unless otherwise noted. FR: females with regular cycles; FI: females with irregular cycles.

| Variable | FR (n=77) | FI (n=59) | Male (n=76) | F / $\chi^2$ | P |
| --- | --- | --- | --- | --- | --- |
| *Sociodemographic* |  |  |  |  |  |
| Age (years) | 15.88 ± 1.00 | 15.86 ± 0.88 | 16.09 ± 1.05 | 1.19 | 0.307 |
| SES score | 30.4 ± 10.4 | 29.0 ± 8.5 | 30.1 ± 10.7 | 0.36 | 0.697 |
| School shift<br>(morning/ full-time) | 26 / 51 | 22 / 37 | 19 / 57 | 2.58<sup>a</sup> | 0.275 |
| *Sleep-wake cycle* |  |  |  |  |  |
| Chronotype — MSFsc (h) | 3.76 ± 1.51 | 3.49 ± 1.39 | 3.95 ± 1.53 | 1.58 | 0.209 |
| Sleep onset (h from midnight) | 22.83 ± 1.32 | 22.73 ± 1.52 | 23.14 ± 1.15 | 1.76 | 0.175 |
| Weekdays |  |  |  |  |  |
| Weekend | 24.33 ± 1.74 | 24.04 ± 1.55 | 24.49 ± 1.57 | 1.31 | 0.271 |
| Wake-up time (h from midnight) | 5.17 ± 0.60 | 5.19 ± 0.81 | 5.46 ± 0.69 | 4.06 | 0.019* |
| Weekdays |  |  |  |  |  |
| Weekend | 8.79 ± 1.86 | 8.54 ± 1.94 | 8.93 ± 1.98 | 0.70 | 0.500 |
| Sleep duration (h) | 6.33 ± 1.31 | 6.46 ± 1.40 | 6.33 ± 1.21 | 0.20 | 0.820 |
| Weekdays |  |  |  |  |  |
| Weekend | 8.47 ± 1.54 | 8.50 ± 1.49 | 8.44 ± 1.43 | 0.03 | 0.969 |
| Sleep debt (h) | 2.13 ± 1.88 | 2.05 ± 1.93 | 2.11 ± 1.71 | 0.04 | 0.963 |
| Social jetlag (h) | 2.56 ± 1.38 | 2.66 ± 2.81 | 2.44 ± 1.44 | 0.23 | 0.794 |
| Sleep quality<br>PSQI (0–21) | 7.43 ± 2.20 | 7.85 ± 2.94 | 7.04 ± 2.97 | 1.48 | 0.229 |
| Daytime sleepiness<br>PDSS (0–32) | 20.42 ± 4.18 | 20.83 ± 4.03 | 18.63 ± 5.06 | 4.83 | 0.009** |
| *Mental health* |  |  |  |  |  |
| Stress<br>DASS-21 (0–42) | 18.99 ± 9.76 | 22.88 ± 9.12 | 14.47 ± 9.80 | 12.91 | <0.001** |
| Anxiety<br>DASS-21 (0–42) | 12.81 ± 8.98 | 17.39 ± 10.77 | 9.29 ± 8.50 | 12.47 | <0.001** |
| Depression<br>DASS-21 (0–42) | 16.05 ± 10.34 | 18.64 ± 9.84 | 14.18 ± 10.45 | 3.15 | 0.045* |

Note. Superscript <sup>a</sup> denotes chi-square statistic. \* $p < 0.05$ . \*\* $p < 0.01$ .

### Mental health severity gradient

We identified a significant group gradient for all three mental health outcomes. For the adjusted linear trend (one-step increase in group order: Male → FR → FI), every additional step was associated with higher scores on all outcomes: stress β/step = 4.33 (95% CI [2.68, 5.98], t = 5.14, p < 0.001), anxiety β/step = 4.08 (95% CI [2.48, 5.67], t = 5.00, p < 0.001), and depression β/step = 2.34 (95% CI [0.59, 4.09], t = 2.62, p = 0.010). Adjusted pairwise contrasts are presented in Table 2 and detailed below and Figure 1 illustrates the gradient for all three outcomes.

**Figure 1.**
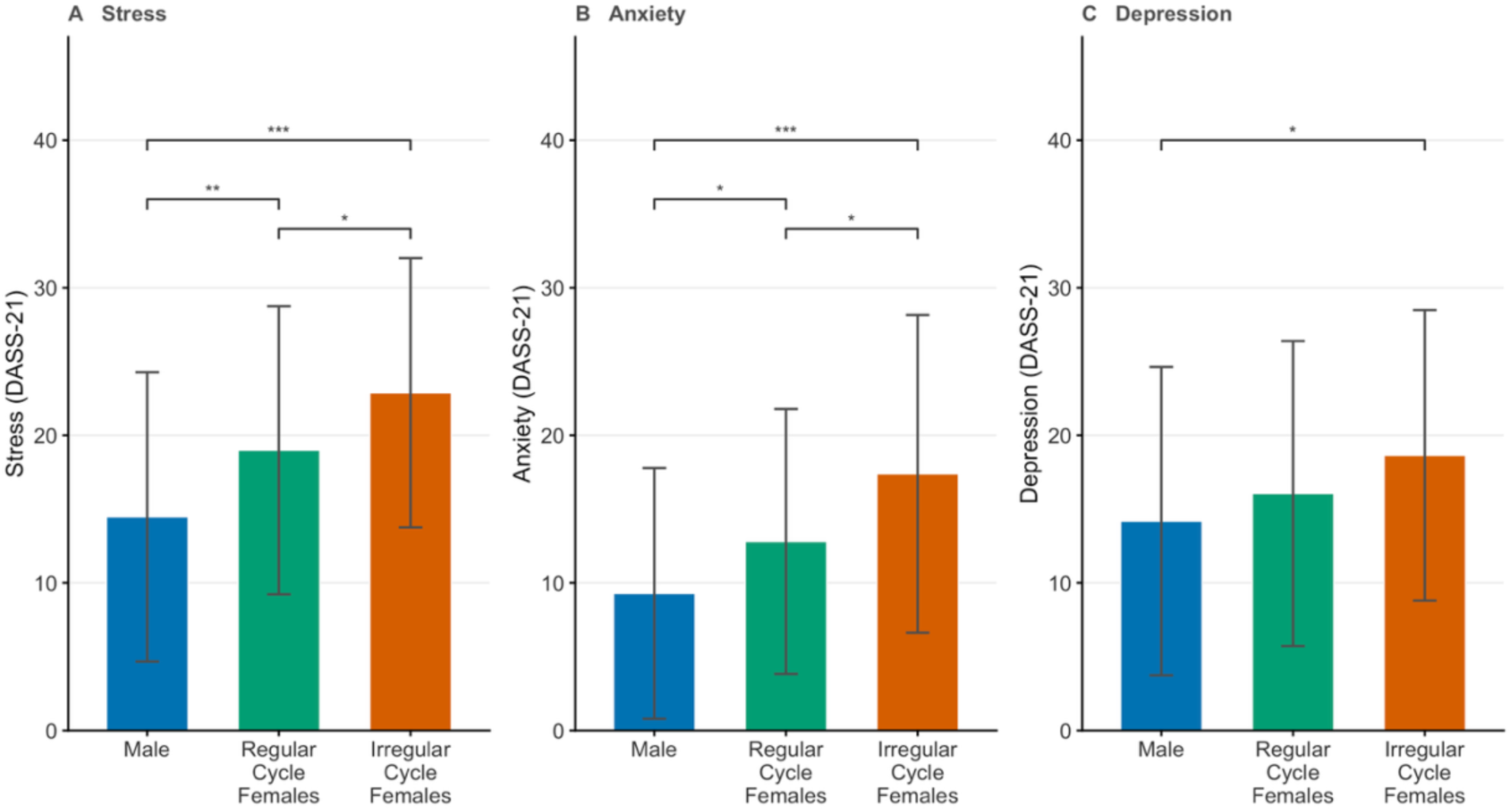
Mean DASS-21 stress, anxiety, and depression scores by group (Male, FR, FI). Error bars = ± 1 SD. Significance brackets reflect FDR-corrected adjusted contrasts from GLM (Table 2). * pᶠᵈᴿ < 0.05, ** pᶠᵈᴿ < 0.01, *** pᶠᵈᴿ < 0.001.

**Table 2.**
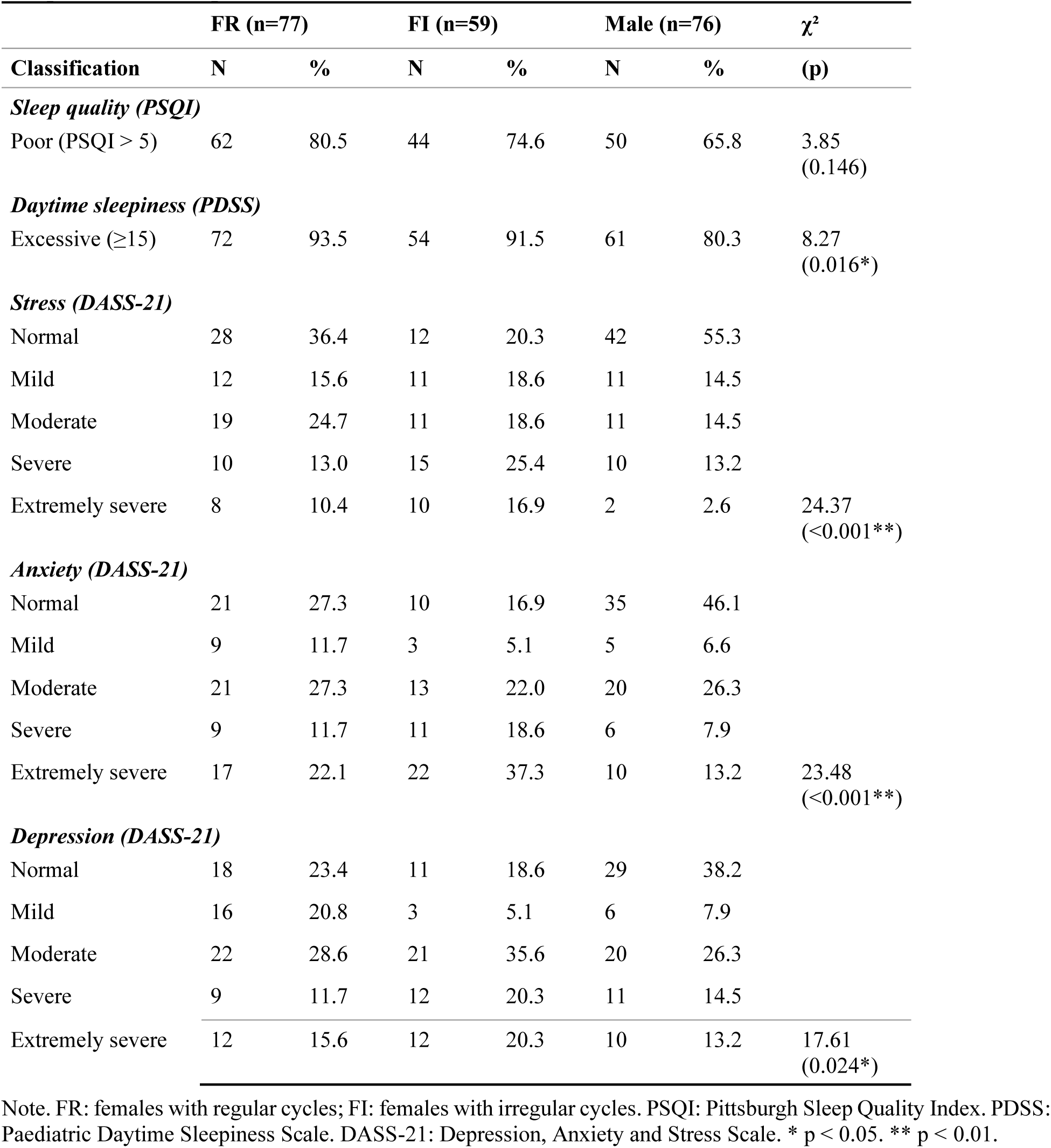
Symptom severity classification proportions (n and %) by group. Chi-square statistic and p-value shown per different outcome.

For stress, the adjusted contrasts exhibit the same monotonic pattern. The FI–Male contrast was the largest (adjusted β = 8.62, 95% CI [5.31, 11.94], p < 0.001, pᶠᵈᴿ < 0.001, d = 0.88). The FR–Male contrast was also significant (adjusted β = 4.71, 95% CI [1.63, 7.79], p = 0.003, pᶠᵈᴿ = 0.009, d = 0.46). Importantly, the FI–FR comparison was also significant after FDR correction (adjusted β = 3.91, 95% CI [0.64, 7.19], p = 0.020, pᶠᵈᴿ = 0.030, d = 0.41). Observed group means were: Male 14.47 ± 9.80, FR 18.99 ± 9.76, FI 22.88 ± 9.12.

For anxiety, the gradient was equally clear: FI scored highest (17.39 ± 10.77), FR intermediate (12.81 ± 8.98), and Male showed the lowest scores (9.29 ± 8.50). The FI–Male contrast exhibited the largest effect size (adjusted β = 8.19, 95% CI [4.97, 11.40], p < 0.001, pᶠᵈᴿ < 0.001, d = 0.85). The FR–Male contrast was also meaningful (adjusted β = 3.69, 95% CI [0.70, 6.67], p = 0.016, pᶠᵈᴿ = 0.024, d = 0.40). The FI–FR contrast was also significant after FDR correction (adjusted β = 4.50, 95% CI [1.32, 7.67], p = 0.006, pᶠᵈᴿ = 0.018, d = 0.47).

Concerning depression symptoms, the FI–Male contrast was significant (adjusted β = 4.69, 95% CI [1.17, 8.21], p = 0.010, pᶠᵈᴿ = 0.010, d = 0.44). However, the FR–Male contrast did not reach statistical significance (β = 2.18, 95% CI [−1.09, 5.45], p = 0.193), and the same pattern was found for the FI–FR contrast after FDR correction (β = 2.51, 95% CI [−0.97, 5.99], p = 0.158, pᶠᵈᴿ = 0.158, d = 0.26), indicating that the depression symptoms gradient is primarily seen through the male–FI contrast rather than a full stepwise progression.

Figure 2 presents symptom severity classification distributions across groups. The gradient in categorical severity was most pronounced for stress and anxiety. Among male participants, 55.3% fell in the Normal stress range and only 2.6% in the Extremely Severe range; among irregular females, only 20.3% were Normal and 16.9% were Extremely Severe. For anxiety, 37.3% of FI scored in the Extremely Severe category, compared to 22.1% of FR and 13.2% of male participants. Omnibus chi-square tests confirmed significant group differences for stress (χ²_(8)_ = 24.37, p = 0.002) and anxiety (χ²_(8)_ = 23.72, p = 0.003). The depression distribution also differed (χ²_(8)_ = 17.61, p = 0.024). However, the depression distribution followed a less pronounced pattern, with 38.2%, 23.4%, and 18.6% in the Normal category and 13.2%, 15.6%, and 20.3% in the Extremely Severe category for males, regular-cycle, and irregular-cycle females respectively.

**Figure 2.**
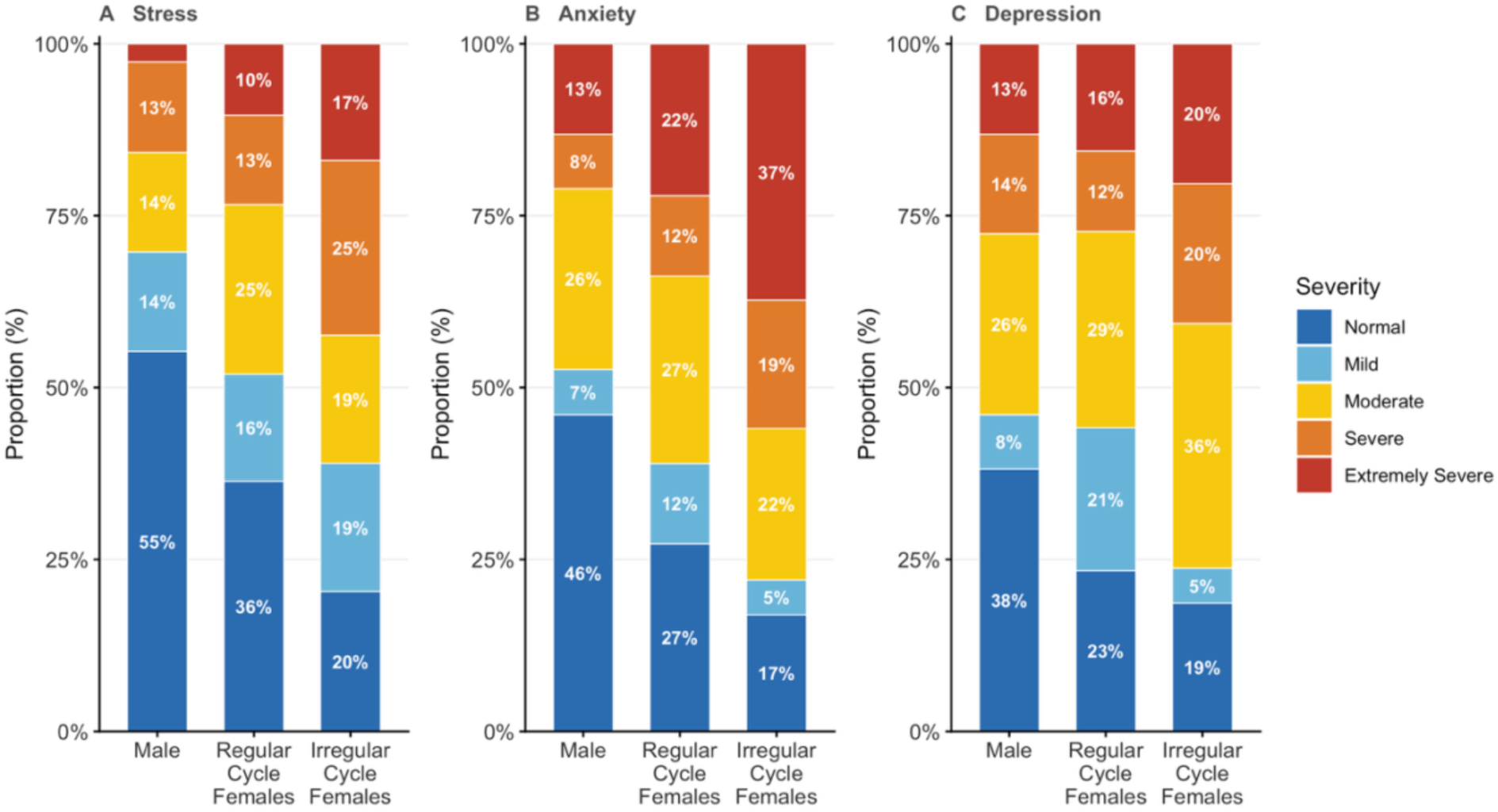
Stacked bar charts of DASS-21 severity classifications (Normal through Extremely Severe) for stress, anxiety, and depression by group. Percentages < 5% are not labelled.

### Socioeconomic status as a sex-specific moderator

The relationship between SES and mental health differed as a function of sex. A Gaussian-identity GLM per outcome demonstrated that the SES slope for male participants was significant and negative for stress (β = −2.68, 95% CI [−4.73, −0.62], p = 0.011) and depression (β = −2.70, 95% CI [−4.89, −0.51], p = 0.017), and marginally significant for anxiety symptoms (β = −2.02, 95% CI [−4.04, 0.00], p = 0.052), which indicates a protective effect of higher SES in male students. The SES × Female interaction term was only statistically significant for stress (β = 3.78, 95% CI [1.12, 6.43], p = 0.006), indicating that the SES-protective effect was substantially weaker in female adolescents. The interaction terms for anxiety (β = 1.50, 95% CI [−1.12, 4.11], p = 0.263) and depression (β = 2.58, 95% CI [−0.26, 5.41], p = 0.076) did not reach statistical significance. The group contrasts remained significant after SES adjustment for stress and anxiety, therefore confirming that the mental health gradient is not explained by socioeconomic differences between groups. Model R² values were 0.157 for stress, 0.134 for anxiety, and 0.079 for depression. When we examined the data by SES quartile, males in the highest quartile (Q4) showed stress scores approximately 6 points lower than those in the lowest quartile (Q1: ∼16.7, Q4: ∼10.4), while the corresponding female values were nearly flat (Q1: ∼19.5, Q4: ∼21.6), consistent with the differential SES buffering observed by sex.

The sleep moderation analysis confirmed that the mental health gradient is independent of sleep parameters. Group × sleep interaction terms were non-significant for all sleep-wake moderators across stress, anxiety and depression (none surviving FDR correction).

## Discussion

In this study tested whether the mental health burden of adolescence follows a graded biological pattern across three groups: male, regular-cycle female and irregular-cycle female adolescents; and whether any such gradient could be attributed to sleep and circadian disruption or moderated by socioeconomic status. Using a cross-sectional design in a Brazilian public-school setting, we assessed depression, anxiety and stress with the DASS-21 and modelled group differences with sex-specific adjusted contrasts, formal sleep-moderation analyses and an SES-by-sex interaction. Three findings emerged. Firstly, symptoms increased monotonically across the groups (male < regular-cycle female < irregular-cycle female), with medium-to-large effects for stress and anxiety and a weaker, male-driven gradient for depression. Secondly, this gradient persisted when sleep and circadian variables were entered as moderators, indicating that it does not depend on the degree of sleep disruption. Finally, higher socioeconomic status was associated with lower symptom burden in male adolescents but not in female adolescents, revealing a sex-based asymmetry in the protective value of socioeconomic resources. Together, these results position menstrual cycle irregularity as a graded biological determinant of adolescent mental health that operates independently of sleep.

A central observation emerging from the present data is that mental health burden was elevated across the entire sample, irrespective of sex or menstrual cycle regularity. Using DASS-21 severity thresholds [21], mean scores for males fell in the Normal range for stress (14.47; threshold ≤ 14), the Mild range for anxiety (9.29; Mild: 8–9), and the Moderate range for depression (14.18; Moderate: 14–20). The regular menstrual cycle group showed Mild stress and Moderate anxiety and depression, while irregular-cycle females showed Moderate stress and depression and borderline Moderate-to-Severe anxiety (17.39; Severe threshold ≥ 20). Considering that no group, not even the male one, fell in the Normal range simultaneously in all three subscales, our results indicate that clinically relevant symptoms of stress, anxiety, and depression are not confined to the highest-burden group but represent a population-level scenario, likely shaped by the combination of overall chronic sleep restriction derived from early school start times [29] and the socioeconomic conditions of Brazilian public school system. Thus, the gradient does not contrast a healthy group against a symptomatic one. Rather, it describes degrees of clinical burden within an overall affected population.

The same optics can be applied to the sleep variables. Despite the absence of significant group differences in sleep, the absolute values across all three groups reveal a picture of uniform disruption in sleep. Mean weekday sleep duration was 6.33 hours for males and regular-cycle females and 6.46 hours for irregular-cycle females, which is approximately 1.5 to 2 hours below the 8 to 10 hours recommended for this age group [30]. Mean PSQI scores exceeded the poor sleep quality threshold of 5 in all three groups (FR = 7.43, FI = 7.85, Male = 7.04), meaning that poor sleep quality was the norm across the entire sample rather than a characteristic of any particular group. Social jetlag exceeded a 2 hour mean in all groups (FR = 2.56 h, FI = 2.66 h, Male = 2.44 h), a level consistently associated with poorer mental health outcomes in adolescents [31]. Mean daytime sleepiness scores were well above the excessive sleepiness threshold of 15 in all three groups (FR = 20.42, FI = 20.83, Male = 18.63), indicating that excessive daytime sleepiness was universal in this sample. Taken together, these levels suggest that the early 7:00–7:30 am school start times in this sociocultural context impose a substantial sleep burden on all students, similar to described in adolescents around the world [29,32], independent of the sex and regularity of the menstrual cycle.

Within this context, this study was specifically designed to test the hypothesis that mental health burden in adolescents follows a graded biological pattern, with males at the lowest end, regular-cycle females in the middle, and irregular-cycle females at the top end. Our data support this hypothesis clearly for stress and anxiety, with medium-to-large effect sizes (R² = 0.16 for stress, 0.13 for anxiety) and significant adjusted contrasts for all pairwise comparisons between groups. For depression the gradient was present in direction and magnitude but the FI–FR contrast did not survive FDR correction (pᶠᵈᴿ = 0.158). This indicates that the depression gradient is primarily driven by the male–FI difference. Beyond replicating the expected direction of group differences, the most noteworthy finding of the present study is that the mental health gradient persists when sleep variables are formally entered as moderators. Group × sleep interaction terms were non-significant for all sleep moderators across stress, anxiety and depression. This systematic null finding should not be considered as a limitation, because it is informative. It suggests that the mental health gradient does not depend on the level of sleep disruption.

The dominant account in the sleep literature positions sleep disruption as the key mediating pathway between sex and mental health outcomes [33,34]. Our findings challenge this idea, but specifically in the context of menstrual irregularity, as the gradient persists regardless of how well, for how long or how regular is sleep in adolescents. Therefore, the mechanisms driving the gradient must operate independently of sleep. We could assume the direct neuroendocrine effects of dysregulated ovarian hormones on stress reactivity and mood regulation as potential mechanistic candidates [4]. The list of candidates would include altered HPA axis sensitivity [35], and somatic symptoms associated with cycle irregularity, such as dysmenorrhea and mood fluctuations [8]. As a natural extension of the present work and to try to disentangle these pathways, longitudinal designs with hormonal measurement would be necessary.

Our secondary finding concerning SES adds an important social dimension with potential implications. The formal SES × Female interaction was statistically significant for stress (β = 3.78, p = 0.006) but not for anxiety or depression (both p > 0.07), with all three in the same direction. The SES slope for males was significantly negative for stress and depression and marginal for anxiety, confirming that higher SES confers measurable mental health protection in male students. This sex-based asymmetry has not been previously reported in an adolescent sample. Higher socioeconomic resources may reduce stress burden through pathways available to male adolescents that are bypassed in females, possibly due to heightened HPA axis sensitivity during pubertal maturation [36,37]. This is consistent with allostatic load theory’s proposition that accumulated biological burden limits the buffering potential of social and economic resources [38–40]. Thus, at least in female adolescents, the fact that the interaction is significant for stress but not for anxiety or depression may reflect greater sensitivity of the stress subscale to socioeconomic context, or insufficient power for the smaller anxiety and depression effects and deserves follow-up in a larger and more socioeconomically diverse sample.

The GLM models explained 15.7% of variance in stress and 13.4% in anxiety (R² values), with large effect sizes (d = 0.88 for FI vs Male stress; d = 0.85 for FI vs Male anxiety), which is clinically meaningful given the population. The proportions of students in the severe and extremely severe anxiety categories (37.3% for the irregular-cycle female versus 13.2% for the male adolescent group) have direct implications for school psychology services and educational wellbeing policies. Early identification of adolescents with irregular menstrual cycles may serve as a low-cost triage tool for elevated anxiety and stress risk, supporting the integration of menstrual health enquiries into school-based mental health routines. While delaying school start times remains an evidence-based strategy for reducing sleep restriction and improving adolescent health [29], our findings indicate that sleep-focused interventions alone may be insufficient. Biological factors specific to female adolescents, including menstrual cycle characteristics, should also be considered when designing school-based health programmes for adolescents.

However, it is important to highlight several limitations in our study. First, the regularity of the menstrual cycle was self-reported and not confirmed by hormonal measurement, introducing a potential classification error. Second, the cross-sectional design precludes causal inference, which makes it plausible that pre-existing mental health problems contribute to menstrual cycle irregularity rather than the reverse. Third, the sample was drawn from a single urban region in Brazil and from schools with atypically early start times relative to international norms, which may limit generalisability to adolescents in different educational systems or socio-cultural contexts.

Fourth, the study did not collect data on dysmenorrhea severity, premenstrual symptoms or body mass index, all of which may confound the irregularity–mental health relationship. Thus, future studies should combine hormonal data, longitudinal assessment of cycle regularity and mental health trajectories while accounting for intraindividual changes across the menstrual cycle, as well as objective sleep measurement, such as actigraphy, to establish the direction and mechanisms of the observed associations.

## Conclusion

This study provides the first evidence that stress and anxiety in adolescents follow a graded biological pattern defined by menstrual cycle regularity. In a sample of 212 students with early school start times, female adolescents with irregular menstrual cycle showed the highest mental health burden, regular-cycle students showed intermediate burden, and males the lowest, a gradient that persisted after covariate adjustment and was not moderated by sleep. Socioeconomic status was protective against mental health burden in males but not in female adolescent students, reinforcing that the vulnerability is biologically based and not responsive to social buffering alone. These findings suggest that menstrual cycle irregularity may serve as a low-cost indicator of elevated mental health risk in adolescent females, supporting its potential integration into school-based health monitoring programmes.

## Acknowledgements

We would like to thank the school’s teaching teams for their collaboration and support, and especially the students who agreed to participate in this research. We would also like to thank the students Izis Lima Ribeiro and Ticyana Dias Costa Silva who helped with data collection and Dr. John Fontenele Araújo, Dr. Maria Bernardete Cordeiro de Sousa and Dr. Fernando Louzada for their valuable suggestions.

## Declaration of Generative AI and AI-assisted technologies in the writing process

During the preparation of this work the authors used Claude (Anthropic, Claude Sonnet 5) in order to refining language, improving clarity, and formatting portions of this manuscript. After using this tool/service, the authors reviewed and edited the content as needed and take full responsibility for the content of the publication.

